# Brief communication: Long-term absence of Langerhans cells alters the gene expression profile of keratinocytes and dendritic epidermal T cells

**DOI:** 10.1101/844530

**Authors:** Qingtai Su, Aurélie Bouteau, Jacob Cardenas, Balaji Uthra, Yuanyaun Wang, Cynthia Smitherman, Jinghua Gu, Botond Z. Igyártó

## Abstract

Tissue-resident and infiltrating immune cells are continuously exposed to molecules derived from the niche cells that often come in form of secreted factors, such as cytokines. These factors are known to impact the immune cells’ biology. However, very little is known about whether the tissue resident immune cells in return also affect the local environment. In this study, with the help of RNA-sequencing, we show for the first time that long-term absence of epidermal resident Langerhans cells (LCs) led to significant gene expression changes in the local keratinocytes and resident dendritic epidermal T cells. Thus, immune cells might play an active role in maintaining tissue homeostasis, which should be taken in consideration at data interpretation.

## INTRODUCTION

The effect of tissue environment on immune cells has been widely studied. Tissue microenvironment through an unknown mechanism is capable of shaping the chromatin landscapes of macrophages, which results in tissue-specific functions of macrophages(1). DC populations in different tissues display tissue-specific diversity and functions(2), and thus, it is anticipated that the close communication between DCs and the tissue microenvironment might endow them with functional diversity and plasticity. It is well documented that keratinocytes for example can regulate immune responses by affecting epidermal resident, antigen presenting Langerhans cells’ biology through secretion of cytokines and other factors(3). Langerhans cells (LCs) are a subset of dendritic cells (DCs) that are radiation-resistant and reside in the epidermis, where they are tightly attached to the surrounding keratinocytes(4). LCs participate in promotion of self-tolerance, anti-fungal immunity, skin immunosurveillance, and protective humoral immune responses(5). In this study, we tested the idea whether long-term absence of an immune cell, LCs from the epithelial environment, affect the constituent KCs and the resident dendritic epidermal T cells (DETCs). Here we show, to our knowledge, first-time evidence that long-term absence of an immune cell can lead to significant changes in the niche cells and to an altered tissue microenvironment.

## MATERIALS AND METHODS

### Mice

huLangerin-DTA (LC^−/−^) mice have been previously described(6). All experiments were performed with 8 weeks old littermate-controlled mice. Mice were housed in microisolator cages and fed autoclaved food. The Baylor Institutional Care and Use Committee approved all mouse protocols.

### Flow cytometry and cell sorting

Single-cell suspensions of flank skin were obtained and stained as previously described(7). Cell suspensions were directly labeled with fluorochrome-conjugated antibodies for cell surface markers anti-MHC-II, anti-CD45 and fixable Viability Dye. KCs (MHC-II^−^, CD45^−^, live events) and DETCs (MHC-II^−^, CD45^+^, live events) were sorted on flow cytometer. Stringent doublets discrimination and live/dead gating were used to exclude possible contaminants and dead cells, respectively.

### RNA preparation

Total RNA was isolated from cell lysates using the RNeasy Micro Kit (Qiagen) including on-column DNase digestion. Total RNA was analyzed for quantity and quality using the RNA 6000 Pico Kit (Agilent).

### Sequencing Library Preparation

Poly-A enriched NGS library construction was performed using the KAPA mRNA Hyper Prep Kit (KAPA Biosystems) using 50ng of input total RNA according to manufacturer’s protocol using 16 amplification cycles. Quality of the individual libraries was assessed using the High Sensitivity DNA Kit (Agilent). Individual libraries were quantitated via qPCR using the KAPA Library Quantification Kit, Universal (KAPA Biosystems) and equimolar pooled. Final pooled libraries were sequenced on an Illumina NextSeq 500 with paired-end 75 base read lengths.

### Bioinformatics analysis

Raw sequencing reads assessed for quality using FASTQC software(8). The adapters were trimmed and low-quality reads (< 20) were filtered using cutadapt(9). Reads were aligned to the mouse reference genome (GRCm38) using hisat2. Aligned SAM files were converted to BAM format using samtools(10) and featureCounts(11) was used to quantify total number of counts for each gene.

### RNA-seq analysis

Transcripts with low expression, i.e., count-per-million (CPM) > 1 in less than two samples, were removed from downstream analysis, leaving 14,964 transcripts. Differential gene expression (DGE) analysis was performed using DESeq2(12) and comparisons were made between LC^−/−^ and WT within DETC and KC cell populations.

### Pathway and Gene Ontology analysis

Two approaches to pathway and Gene Ontology (GO) analysis were used(13). The Database for Annotation, Visualization and Integrated Discovery (DAVID)(14) was used for functional annotation of significantly regulated genes based on false discovery rate (FDR) < .05 and fold change (FC) cut-off of 1.5 for each comparison. Additionally, a fast implementation of pre-ranked Gene Set Enrichment Analysis (FGSEA) using the fgsea R package(15,16) was performed on KEGG and GO gene sets obtained from the Molecular Signatures Database v6.2 (MSigDB)(17).

### RNA-seq data visualization

Counts were normalized using the median-of-ratios method(18) and log2 transformed for data visualization. Principal component analysis (PCA) and hierarchical clustering were performed using the R. The transcripts of all heatmaps were hierarchically clustered using Euclidean distance and complete linkage function. Heatmaps were plotted using the NMF package(19), while PCA and volcano plots were made using ggplot2(20).

## RESULTS

### Long-term absence of LCs leads to gene expression changes in KCs and DETCs

To determine the possible effect of the absence of LCs on the cells of the epidermis, we took advantage of the huLangerin-DTA mice (hereafter LC^−/−^), which lack LCs starting from birth(6). Thus, for these mice, KCs and DETCs develop, differentiate, and function in the absence of mature LCs. Epidermal cells suspensions were generated from a cohort of LC^−/−^ mice, along with littermate WT controls (**Figure 1a**). After staining with specific markers, the KCs and DETCs were sorted using flow cytometer, and RNA-sequencing performed. Unsupervised PCA of the expression data revealed that KCs and DETCs, which developed in the absence of LCs, clearly clustered away from their WT counterparts (**Figure 1b**). We identified 1220 up- and 537 downregulated genes in KCs, while in DETCs, we identified 880 up- and 214 downregulated genes using a false discovery rate (FDR) <0.05 (**Figure 1c**). Out of the upregulated genes, 348 (19.9%) were common between KCs and DETCs, while 22 genes (3.02%) were commonly downregulated (**Figure 1c**). Next, we performed hierarchical clustering of differentially expressed genes with at least 2-fold change and plotted heatmaps to show the distinct patterns of up- and downregulated genes in KCs and DETCs (**Figure 1d**). We used color-coded volcano plots to better capture and visualize the common and cell specific changes in gene expression (**Figure 2a**). We observed that nerve growth factor (NGF) was highly upregulated in KCs and DETCs. NGF is part of the neurotrophin family and is involved in the differentiation and survival of neural cells(21), which suggest that LCs might directly or indirectly regulate nerve homeostasis in the epidermis.

**Figure 1.**
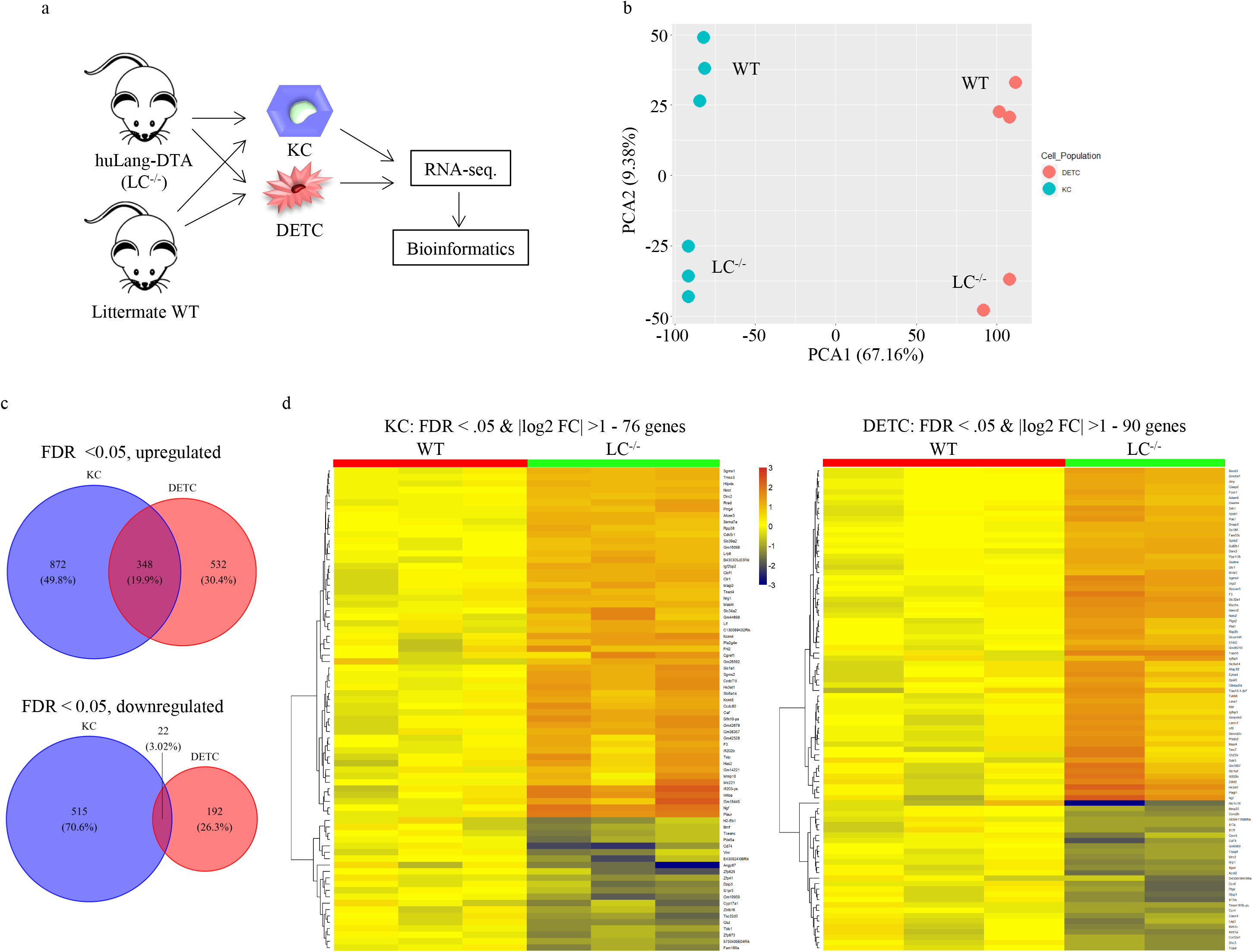
Absence of LCs leads to gene expression changes in KCs and DETCs. **a**. Experimental flow. KCs and DETCs were flow sorted from LC-deficient (LC^−/−^) and littermate WT controls and RNA-seq. performed. The resulting data were then subjected to bioinformatic analyses. **b**. Principal component analysis of the RNA-seq. data. Each dot represents a separate animal. **c**. The overlaps between the genes that were up- (top) or downregulated (bottom) in the absence of LCs in KCs and DETCs are presented in forms of Venn diagrams; FDR<0.05. **d**. Heatmap presentation of the genes that showed two-fold changes between LC^−/−^ and WT mice. KCs (left) and DETC (right). FDR<0.05.

**Figure 2.**
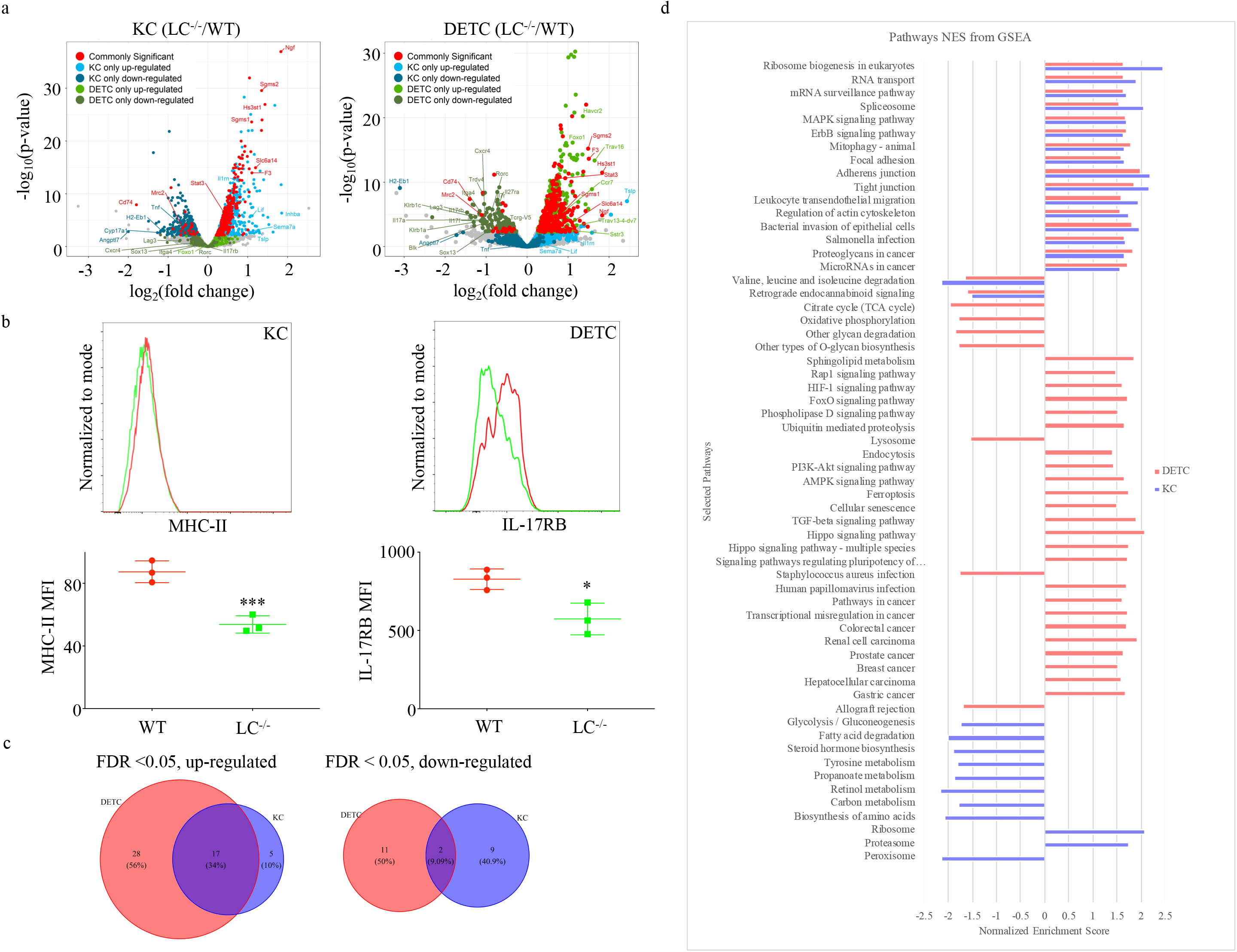
LCs have common and cell specific effects on KCs’ and DETCs’ biology. **a**. Common and cell specific gene expression changes are presented in form of color-coded volcano plots. KCs (left) and DETCs (right). **b**. Flow cytometry confirmation of the RNA-seq. data on protein levels. Each dot represents a separate mouse. Two tailed Student’s t-test. *p<0.05, ***p<0.001. **c**. The overlap between up- or downregulated regulated KEGG pathways in KCs and DETCs from LC^−/−^ mice are presented in forms of Venn diagrams. **d**. Selected KEGG pathways altered in KCs and DETCs in the absence of LCs are depicted based on normalized enrichment scores (NES). FDR<0.05.

*TSLP* was specifically upregulated in the KCs in the absence of LCs, which is in concordance with a recently published article by Lee et al.(22) TSLP is a known regulator of the Th2 responses and it is also needed for mast cell homeostasis(23,24). Dysregulated TSLP production by KCs, in the absence of LCs, could have contributed to altered IgE levels(25) and increased mast cell numbers (unpublished observation on FVB background) observed in the LC^−/−^ mice. The KCs, among others, showed downregulation of the MHC-II pathway genes H2-Eb1 (**Figure 2a and b**), CD74 (invariant chain; also downregulated in DETCs;), and Cyp17a1, a member of the P450 cytochrome family involved in carcinogen metabolism(26,27).

### Lack of LCs might affect DETCs’ biology and homeostasis

We discovered that DETCs downregulated the Th17 pathway associated molecules, including *RORc* transcription factor, *IL-17RB* receptor, and *IL-17A* and *IL-17F* cytokines (**Figure 2a and b**). The IL-17 pathway plays an important role in the DETC’s innate immune function to fight bacterial infections(28). More interestingly, we observed that DETCs showed lower expression of the γ/δ TCRs (*Trdv4* and *Tcrg-V5*) and upregulation of TCR alpha chains (*Trav16* and *Trav13-4-dv7*). Transcription factors and other molecules that regulate the development, differentiation, and homeostasis of DETCs, such as *Sox13, Blk*, and *Il-7r*(29), were also downregulated. Thus, unlike previously published data(30), these data suggest that LCs might directly or indirectly regulate DETCs’ biology and homeostasis, and could contribute to maintain their identity/fitness in the epidermis/periphery in the absence of the thymic environment.

### The absence of LCs affected a variety of different cellular pathways in KCs and DETCs

To gain a broader picture about the effect of the absence of LCs on KCs and DETCs, we performed KEGG pathway analysis on the expression data. We present data of significantly altered pathways using FDR < 0.05. We observed significant overlap of pathways upregulated by KCs and DETCs, but very minimal overlap of downregulated pathways (**Figure 2c**). The commonly upregulated pathways included different forms of cell adhesions (focal, adherent and tight)-, ribosome and RNA biogenesis-, autophagy-, bacterial invasion/infection-, MAPK- and ErbB signaling pathways (**Figure 2d**). Alterations in adhesion molecules and the ErbB signaling pathway in KCs, in the absence of LCs, were recently reported(22,31). The downregulated pathways showed considerably less overlap between these two cell types and included some of the amino acid degradation pathways (**Figure 2d**). KCs showed distinct dysregulation (mostly downregulation) of various metabolic pathways (sugar, protein, fatty acids, hormones, drug, xenobiotics etc.), while DETCs presented with alterations in TGF-β–, Hippo-, oxidative phosphorylation-, citrate cycle-, lipid metabolism-, Staphylococcus aureus infection- etc. pathways (**Figure 2d**). Thus, these data suggest that LCs might have common and cell-specific effects on KCs’ and DETCs’ biology.

## DISCUSSION

Here we bring experimental evidence that long-term absence of LCs leads to gene expression changes in KCs and DETCs. The significant changes discovered by pathway analysis also suggest that KCs’ and DETCs’ biology and hemostasis are likely affected. Further studies are needed to confirm the observed changes and their consequences. It will also be important to determine which LC-derived factors play role in the epidermal homeostasis. Our preliminary IPA Upstream Regulator Analysis identified a list of potential regulators, including cytokines, growth factors, and enzymes (data not shown), known or anticipated to be produced by LCs.

To our knowledge we show for the first time that long-term absence of an immune cell can lead to significant changes in the niche cells and to altered tissue environment. The effect of niche cells on resident immune cells is very much appreciated by the immunologist, however, our findings support the idea that the resident immune cells are not mere passive receivers, but rather play an active and indispensable role in maintaining tissue homeostasis. Thus, studies using constitutive immune-cell knockouts, including LC^−/−^ mice, in which the immunological changes and outcomes were directly attributed to the absence of a specific immune cell, might have to be reassessed(5,6,32–34).

## ACKNOWLEDGEMENTS

We thank the animal facility, flow core and genomics core for their help and support. The Baylor Scott & White Health Foundation supported this work.

## REFERENCES

1. Lavin Y, Winter D, Blecher-Gonen R, David E, Keren-Shaul H, Merad M, et al. Tissue-Resident Macrophage Enhancer Landscapes Are Shaped by the Local Microenvironment. Cell. 2014 Dec;159(6):1312–26.

2. Pakalniškytė D, Schraml BU. Tissue-Specific Diversity and Functions of Conventional Dendritic Cells. In: Advances in Immunology. 2017.

3. Clayton K, Vallejo AF, Davies J, Sirvent S, Polak ME. Langerhans cells-programmed by the epidermis. Frontiers in Immunology. 2017.

4. Romani N, Holzmann S, Tripp CH, Koch F, Stoitzner P. Langerhans cells - dendritic cells of the epidermis. APMIS. 2003 Jul;111(7–8):725–40.

5. Kaplan DH. Ontogeny and function of murine epidermal Langerhans cells. Nat Immunol. 2017 Sep(18):10–1068.

6. Kaplan DH, Jenison MC, Saeland S, Shlomchik WD, Shlomchik MJ. Epidermal langerhans cell-deficient mice develop enhanced contact hypersensitivity. Immunity. 2005 Dec(23):6–611.

7. Yao C, Zurawski SM, Jarrett ES, Chicoine B, Crabtree J, Peterson EJ, et al. Skin dendritic cells induce follicular helper T cells and protective humoral immune responses. J Allergy Clin Immunol. 2015 May;

8. Andrews S. FASTQC A Quality Control tool for High Throughput Sequence Data. Babraham Inst. 2015;

9. Martin M. Cutadapt removes adapter sequences from high-throughput sequencing reads. EMBnet.journal. 2011;

10. Li H, Handsaker B, Wysoker A, Fennell T, Ruan J, Homer N, et al. The Sequence Alignment/Map format and SAMtools. Bioinformatics. 2009;

11. Liao Y, Smyth GK, Shi W. FeatureCounts: An efficient general purpose program for assigning sequence reads to genomic features. Bioinformatics. 2014;

12. Love MI, Huber W, Anders S. Moderated estimation of fold change and dispersion for RNA-seq data with DESeq2. Genome Biol. 2014;

13. Ashburner M, Ball CA, Blake JA, Botstein D, Butler H, Cherry JM, et al. Gene ontology: Tool for the unification of biology. Nature Genetics. 2000.

14. Huang DW, Sherman BT, Lempicki RA. Systematic and integrative analysis of large gene lists using DAVID bioinformatics resources. Nat Protoc. 2009;

15. Sergushichev AA. An algorithm for fast preranked gene set enrichment analysis using cumulative statistic calculation. bioRxiv. 2016;

16. Subramanian A, Tamayo P, Mootha VK, Mukherjee S, Ebert BL, Gillette MA, et al. Gene set enrichment analysis: A knowledge-based approach for interpreting genome-wide expression profiles. Proc Natl Acad Sci U S A. 2005;

17. Liberzon A, Subramanian A, Pinchback R, Thorvaldsdóttir H, Tamayo P, Mesirov JP. Molecular signatures database (MSigDB) 3.0. Bioinformatics. 2011;

18. Anders S, Huber W. Differential expression analysis for sequence count data. Genome Biol. 2010;

19. Gaujoux R, Seoighe C. A flexible R package for nonnegative matrix factorization. BMC Bioinformatics. 2010;

20. Ginestet C. ggplot2: Elegant Graphics for Data Analysis. J R Stat Soc Ser A (Statistics Soc. 2011;

21. Skaper SD. Nerve growth factor: a neuroimmune crosstalk mediator for all seasons. Immunology. 2017.

22. Lee HJ, Kim TG, Kim SH, Park JY, Lee M, Lee JW, et al. Epidermal Barrier Function Is Impaired in Langerhans Cell-Depleted Mice. J Invest Dermatol. 2019;

23. Han NR, Oh HA, Nam SY, Moon PD, Kim DW, Kim HM, et al. TSLP induces mast cell development and aggravates allergic reactions through the activation of MDM2 and STAT6. J Invest Dermatol. 2014;

24. Leyva-Castillo JM, Hener P, Michea P, Karasuyama H, Chan S, Soumelis V, et al. Skin thymic stromal lymphopoietin initiates Th2 responses through an orchestrated immune cascade. Nat Commun. 2013;

25. Nakajima S, Igyártó BZ, Honda T, Egawa G, Otsuka A, Hara-Chikuma M, et al. Langerhans cells are critical in epicutaneous sensitization with protein antigen via thymic stromal lymphopoietin receptor signaling. J Allergy Clin Immunol. 2012 Apr;129(4):1048––55.e6.

26. Lewis JM, Bürgler CD, Freudzon M, Golubets K, Gibson JF, Filler RB, et al. Langerhans Cells Facilitate UVB-Induced Epidermal Carcinogenesis. J Invest Dermatol. 2015;

27. Modi BG, Neustadter J, Binda E, Lewis J, Filler RB, Roberts SJ, et al. Langerhans cells facilitate epithelial DNA damage and squamous cell carcinoma. Science (80-). 2012;

28. Cho JS, Pietras EM, Garcia NC, Ramos RI, Farzam DM, Monroe HR, et al. IL-17 is essential for host defense against cutaneous Staphylococcus aureus infection in mice. J Clin Invest. 2010;

29. Kang J, Malhotra N. Transcription Factor Networks Directing the Development, Function, and Evolution of Innate Lymphoid Effectors. Annu Rev Immunol. 2015;

30. Taveirne S, De Colvenaer V, Van Den Broeck T, Van Ammel E, Bennett CL, Taghon T, et al. Langerhans cells are not required for epidermal V 3 T cell homeostasis and function. J Leukoc Biol. 2011;

31. Shipman WD, Chyou S, Ramanathan A, Izmirly PM, Sharma S, Pannellini T, et al. A protective Langerhans cell keratinocyte axis that is dysfunctional in photosensitivity. Sci Transl Med. 2018;

32. Igyártó BZ, Haley K, Ortner D, Bobr A, Gerami-Nejad M, Edelson BT, et al. Skin-resident murine dendritic cell subsets promote distinct and opposing antigen-specific T helper cell responses. Immunity. 2011 Aug(35):2–260.

33. Igyártó BZ, Jenison MC, Dudda JC, Roers A, Müller W, Koni PA, et al. Langerhans cells suppress contact hypersensitivity responses via cognate CD4 interaction and langerhans cell-derived IL-10. J Immunol. 2009 Oct(183):8–5085.

34. Obhrai JS, Oberbarnscheidt M, Zhang N, Mueller DL, Shlomchik WD, Lakkis FG, et al. Langerhans cells are not required for efficient skin graft rejection. J Invest Dermatol. 2008;

